# Gene regulatory network inference in soybean upon infection by *Phytophthora sojae*

**DOI:** 10.1101/2022.10.19.512983

**Authors:** Brett Hale, Sandaruwan Ratnayake, Ashley Flory, Ravindu Wijeratne, Clarice Schmidt, Alison E. Robertson, Asela J. Wijeratne

**Affiliations:** Molecular Biosciences Graduate Program, Arkansas State University, State University, AR, USA; Arkansas Biosciences Institute, Arkansas State University, State University, AR, USA; College of Science and Mathematics, Arkansas State University, State University, AR, USA; Houston High School, Germantown, TN, USA; Department of Plant Pathology and Microbiology, Iowa State University, Ames, IA, USA

**Keywords:** soybean, *P. sojae*, transcription factor, gene regulation, gene regulatory network, deep learning

## Abstract

*Phytophthora sojae* is a soil-borne oomycete and the causal agent of Phytophthora root and stem rot (PRR) in soybean (*Glycine max* [L.] Merrill). Yield losses attributed to *P. sojae* are devastating in disease-conducive environments, with global estimates surpassing 1.1 million tonnes annually. Historically, management of PRR has entailed host genetic resistance (both vertical and horizontal) complemented by disease-suppressive cultural practices (e.g., oomicide application). However, the vast expansion of complex and/or diverse *P. sojae* pathotypes necessitates developing novel technologies to attenuate PRR in field environments. Therefore, the objective of the present study was to couple high-throughput sequencing data and deep learning to elucidate molecular features in soybean following infection by *P. sojae*. In doing so, we generated transcriptomes to identify differentially expressed genes (DEGs) during compatible and incompatible interactions with *P. sojae* and a mock inoculation. The expression data were then used to select two defense-related transcription factors (TFs) belonging to WRKY and RAV families. DNA Affinity Purification and sequencing (DAP-seq) data were obtained for each TF, providing putative DNA binding sites in the soybean genome. These bound sites were used to train Deep Neural Networks with convolutional and recurrent layers to predict new target sites of WRKY and RAV family members in the DEG set. Moreover, we leveraged publicly available Arabidopsis (*Arabidopsis thaliana*) DAP-seq data for five TF families enriched in our transcriptome analysis to train similar models. These Arabidopsis data-based models were used for cross-species TF binding site prediction on soybean. Finally, we created a gene regulatory network depicting TF-target gene interactions that orchestrate an immune response against *P. sojae*. Information herein provides novel insight into molecular plant-pathogen interaction and may prove useful in developing soybean cultivars with more durable resistance to *P. sojae*.

**Author Summary:** Global food security is threatened continually by plant pathogens. One approach to circumvent these disease-causing agents entails understanding how hosts balance primary growth and defense upon pathogen perception. Molecular signatures of perception-rendered defense may be leveraged subsequently to develop resistant/tolerant crop plants. Additionally, evidence suggests that the plant immune system is characterized by tuning primary and secondary metabolic activity via transcription factor-mediated transcriptional reprogramming. Therefore, we investigated transcription factor-target gene interactions in soybean upon infection by compatible and incompatible races of *Phytophthora sojae*. Through transcriptome analysis, we found that the interactions elicited vast, overlapping transcriptional responses and identified overrepresented, defense-related transcription factor families. We then generated/acquired DNA-protein interactome data for the most represented transcription factor families in the transcriptome analysis and trained deep learning-based models to predict novel transcription factor targets. Transcription factor/target gene metrics were used to construct a gene regulatory network with prioritized components. We identified hub transcription factors belonging to WRKY and ERF families, the majority of which function in response to various biotic and abiotic stressors. These findings propose novel regulators in the soybean defense response to *Phytophthora sojae* and provide an avenue for the investigation of transcription factor-target gene interactions in plants.

## Introduction

*Phytophthora sojae* Kaufmann and Gerdemann is a hemibiotrophic, homothallic oomycete that renders significant yield losses in soybean (*Glycine max* [L.] Merrill). The pathogen can infect host plants at any developmental stage and is denoted by damping-off in seedlings (early season) as well as root rot and subsequent chlorosis/necrosis in aboveground tissue (late season) [1]. In addition, *P. sojae* oospores can persist in a production environment for several years, limiting the efficacy of most cultural management strategies [2,3]. Thus, the most economical and environmentally benign method to manage the pathogen is the deployment of horizontal and/or vertical host genetic resistance [4]. Horizontal resistance is quantitatively inherited and provides some level of protection against all *P. sojae* pathotypes. However, early-season efficiency is reliant upon complementation with cultural practices [3] as it is only active after the first true leaf stage. Moreover, the polygenic character of horizontal resistance hinders introgression into germplasm [5,6]. Alternatively, vertical resistance (i.e., incompatibility) embodies the classic gene-for-gene concept and renders complete protection against specific pathotypes in a monogenic manner [7]. The selective pressures imposed by vertically resistant soybean have increased the virulence profile of *P. sojae* populations, restricting the use of cultivars with a specific *Resistance to P. sojae* (*Rps*) gene to 8-15 years [2,8]. Therefore, a deeper understanding of the molecular mechanisms governing soybean defense against *P. sojae* is needed to overcome pathogen evolution and ultimately attenuate disease.

During a compatible (virulent) soybean-*P. sojae* interaction, the host plant perceives microbe-associated molecular patterns (MAMPs)/pathogen-associated molecular patterns (PAMPs) and elicits PAMP-triggered immunity (PTI), a basal immune response effective against non-adapted pathogens [9]. Conversely, *P. sojae* secretes *Avirulence* (*Avr*) gene-encoded effector proteins that suppress components of PTI and promote disease. During incompatibility, a receptor encoded by an *Rps* gene recognizes the cognate *Avr* gene product and activates effector-triggered immunity (ETI), a hypersensitive immune response that potentiates PTI and confers resistance to *P. sojae* [7,9,10]. The combined efforts of PTI and ETI to mitigate disease during incompatibility exemplify the zig-zag model of Jones and Dangl [11] and portray PTI and ETI as distinct events that occur consecutively. A growing body of evidence obscures these boundaries, particularly in plant-*Phytophthora* interactions, instead suggesting that plant defense spans a PTI:ETI continuum [12,13]. For these reasons, Wang et al. [14] proposed a three-layered model of plant immunity comprising a recognition layer, a signal-integration layer, and a defense-action layer. In the context of soybean-*P. sojae* interaction, our understanding of the signal-integration layer remains the most fragmented.

Intra- and/or extracellular pathogen perception triggers a dynamic, highly sophisticated signaling network that balances primary and secondary metabolic activity in a manner preservative of host fitness [15]. Signal integration and convergence accompanying this coordinated stress response are mediated by transcription factors (TFs) and transcriptional cofactors that comprise sensory regulatory networks embedded within phytohormone signaling pathways [16,17]. Dynamism and amplification of such networks are determined by physical interaction between TFs and nucleocytoplasmic receptors [18,19], TF phosphorylation by 2+ mitogen-activated protein kinase (MAPK) cascades [20], and feedback regulation of Ca^2+^ signaling, among other mechanisms [17]. While the abundance and diversity of TFs required for immunity vary across plant species and pathosystems [21], elucidated sensory regulatory networks tend to possess members of the bHLH, bZIP, ERF, MYB, NAC, and WRKY families [17,22,23] that collectively direct transcriptional reprogramming of downstream target genes [16]. Isolated studies have evidenced transcriptional reprogramming in soybean upon infection by *P. sojae* [24,25,26,27] and have identified various TFs associated with defense [28,29,30,31,32,33,34,35,36,37]; yet mechanistic insight regarding TF-target gene interactions and their organization within larger hierarchical networks is lacking. This systems-level information can be unraveled using gene regulatory networks (GRNs) [38], and regulatory hubs identified subsequently through analyses of network tunability and redundancy [21].

In a simplistic model of gene regulation, TFs bind to regulatory DNA motifs in target genes to modulate transcriptional activity [39]. GRNs can be used to discern static and spatiotemporal interactions between TFs and DNA motifs as well as interaction abundance, topology, and influence on target gene expression [40,41]. Various experimental and computational methods have been developed to study gene regulation and function for a phenomenon of interest (e.g., disease resistance) [38]. As an example, TF-DNA interactome approaches such as chromatin immunoprecipitation followed by sequencing (ChIP-seq) and DNA-affinity purification and sequencing (DAP-seq) allow the identification of many TF binding sites at once and can be used to validate interactions inferred by gene expression analysis [41,42]. However, these methodologies are technically- and economically-demanding and are thus difficult to deploy at genome-scale.

Contrarily, *in silico* exploration of TF-target interactions is easily scalable. The majority of such methods leverage guilt-by-association approaches that cannot necessarily predict causality and are thus limited in terms of elucidating regulatory pathways [43]. One can overcome this by using a bottom-up approach to identify *cis*-regulatory elements (CREs), which modulate gene expression by recruiting TFs, as a means to presume TF-target gene interactions. The most popular approach to find CREs is to employ a supervised motif method using a position-specific score matrix (i.e., position weight matrix) and map CREs to a promoter [44]. However, given that CREs are often degenerate and short, this method suffers from high false positive rates. Improvement can be made by considering the evolutionary conservation of CREs (albeit all functional CREs are not necessarily evolutionary conserved) or gene co-regulation. More recently, Deep Neural Network (DNN)-based methods were developed to detect TF binding sites (TFBS) [45,46]. The DNN-based techniques are deemed superior to others given their ability to accept minor CRE variation and sequence context surrounding TFBS and thus transfer across species [47]. For instance, leveraging Arabidopsis (*Arabidopsis thaliana*) cistrome datasets, Akagi et al. [48] constructed convolutional neural network (CNN)-based DNN models for cross-species prediction of TFBS in tomato (*Solanum lycopersicum*). Likewise, Bang et al. [49] used CNN models to predict TFBS in both maize (*Zea mays*) and soybean using maize DAP-seq data. Although high false positive rates were observed by cross-species prediction in the latter study [49], these advances demonstrate the value of DNN-based methods for TFBS prediction within and across plant species.

In the present study, we coupled transcriptomics, *in vitro* TF-DNA interaction profiling, and deep learning to construct a GRN underlying the soybean defense response to *P. sojae* infection (**Fig 1**). We first inoculated hypocotyls of soybean variety Williams 82 (possesses *Rps1k*) with mycelial slurries from *P. sojae* Races 1 or 25, rendering incompatible and compatible interactions, respectively. Transcriptomes were generated from the hypocotyls, and differential gene expression analysis was performed with expression profiles from a mock inoculation serving as a baseline. Following RNA capture-based validation of the experimental design, we assessed TF representation in the differentially expressed gene (DEG) set. The biological significance of overrepresented TF families was inferred by clustering DEGs, assigning functional annotations to each cluster, and observing TF family representation across defense-related clusters. Next, using DAP-seq data for WRKY and RAV TFs differentially expressed in our DEG set, we obtained promoter-localized TFBS to train DNN models for the prediction of novel WRKY and RAV targets. Furthermore, cross-species target prediction was performed for MYB, WRKY, NAC, ERF, and bHLH TF families using DNN models trained with available Arabidopsis DAP-seq data. We observed the representation of predicted targets in our DEG set and used highly confident TF-target predictions to reconstruct a GRN. Findings in this study provide insight into the regulatory mechanisms governing defense against *P. sojae* and provide new/novel avenues for the molecular breeding of soybean.

**Fig 1.**
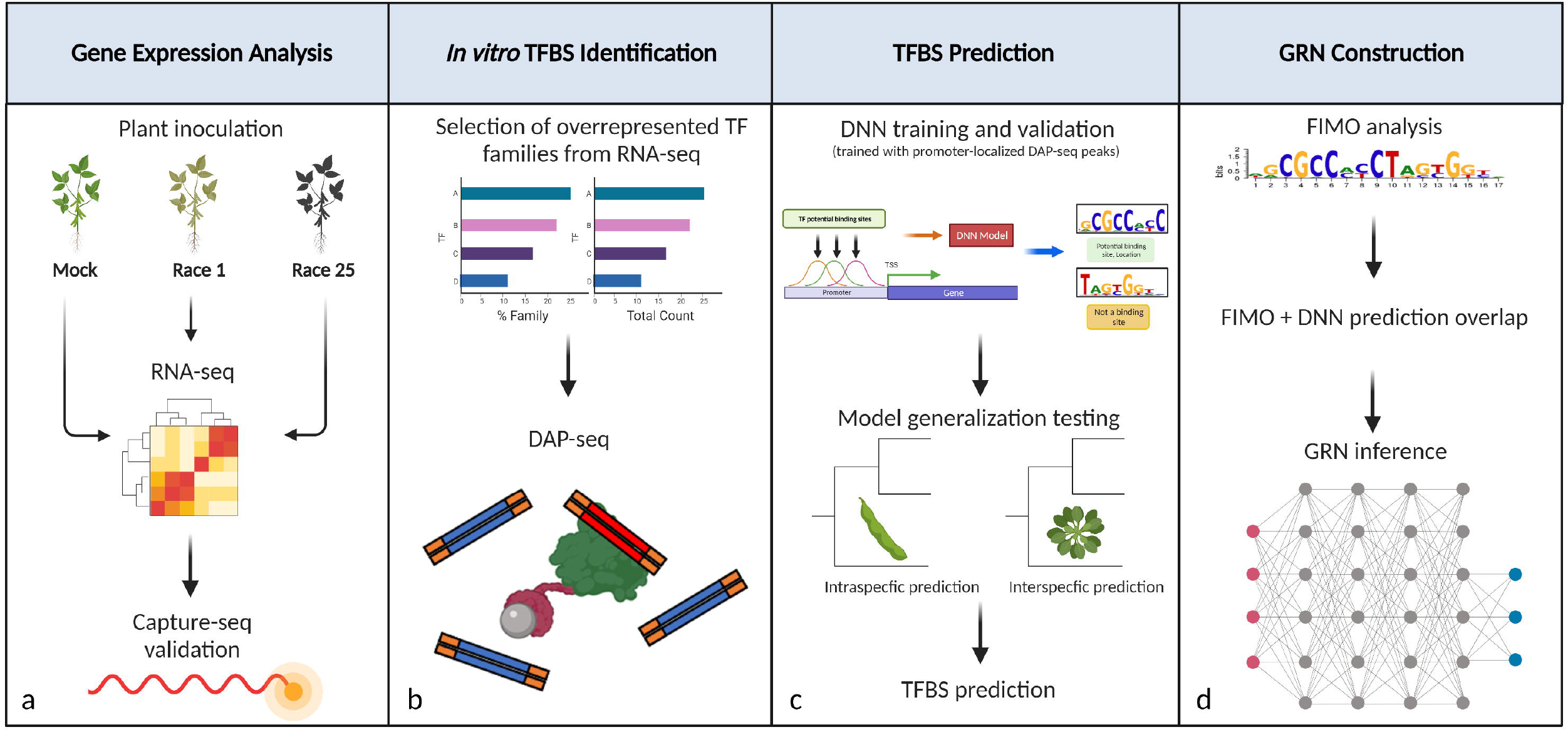
Schematic overview of the study design. (**a**) Soybean plants harboring *Rps1k* were inoculated with a Race 1 *P. sojae* isolate, Race 25 isolate, or sterile media. Inoculated hypocotyls were used for RNA-seq. Capture-seq was performed subsequently to validate the RNA-seq data. (**b**) Overrepresented TF families were identified from the RNA-seq analysis. DAP-seq data was generated/obtained for the families most represented by total abundance and percentage of genome-wide proportion. (**c**) DL models were trained using DAP-seq binding site data. The capacity of some models to generalize across a given TF family was performed intra- and interspecifically. For several TF families of interest, soybean- or Arabidopsis-based DNNs were trained and used to predict TFBS. (**d**) DNN predictions were overlapped with FIMO motif scans, and the highly confident targets were used to construct a GRN.

## Results

### Transcriptome analysis and virulence screening

To identify candidate genes involved in the defense response against *P. sojae*, seeds of soybean variety Williams 82 were grown in germination paper and seedling hypocotyls inoculated with a mycelial slurry from a Race 1 isolate (R1; incompatible interaction), a Race 25 isolate (R25; compatible interaction), or sterile media (Mock) following the procedure of Dorrance et al. [50]. At 24 hrs post-infection (hpi), the hypocotyls were collected and used to generate 13 RNA-seq libraries (4 Mock, 4 R1, and 5 R25) spanning two independent inoculations, RNA isolations, and sequencing runs. Additional seedlings were maintained seven days post-infection (dpi) to compare disease development across treatments (**Fig 2a**) [50]. Collectively, the RNA-seq samples comprised over 560 million 100-bp paired-end reads with a mean mapping rate of 95% (**Table S1**). Principal Component Analysis demonstrated that samples clustered according to treatment and sequencing event. To circumvent the latter, we used ComBat-seq [51], a negative binomial regression model, to correct batch effects. We then performed differential gene expression analysis and removed any genes with expression that had been significantly changed (False Discovery Rate < 0.05) between the two batches. Furthermore, we checked the expression of six genes that displayed stable expression in soybean upon various biotic stresses [52] and a gene possessing a *P. sojae*-inducible promoter (*GmaPPO12*) [53]. The defined reference genes showed no significant differences among treatments in the present study, while *GmaPPO12* showed strong induction due to *P. sojae* infection (**Data S1**). Cooperatively, all R1-inoculated hypocotyls displayed a localized hypersensitive response, while those inoculated with R25 were demarcated by expansive necrotic lesions (**Fig 2a**).

**Fig 2.**
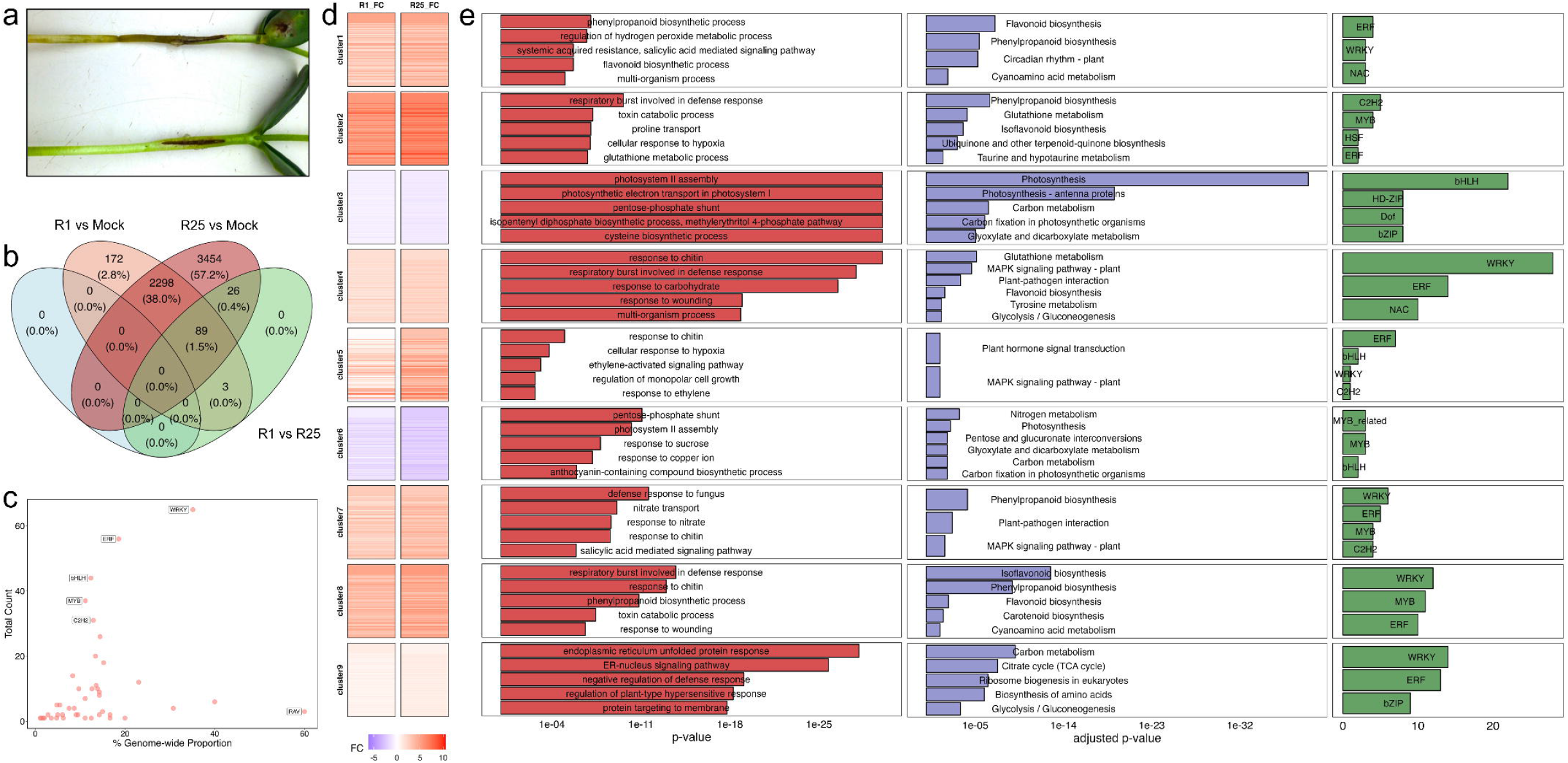
Pathogenicity testing and transcriptome analysis. (**a**) Disease development in Race 25- (top) and Race 1-treated (bottom) hypocotyls at seven days post-infection. (**b**) Venn diagram of DEGs between different treatments. (**c**) TF representation among DEGs from RNA-seq. WRKY was the most represented TF family by total abundance and RAV by the percentage of genome-wide proportion. (**d**) K-means clustering of DEGs. DEGs were assigned to nine co-expression clusters. Of these, seven displayed increased expression (log_2_FC [FC] >0) in infected vs Mock treatments, while two demonstrated decreased expression (FC <0). (**e**) Functional enrichment and TF representation for gene co-expression clusters. (left panel) Top five GO categories by *p*-value. (middle panel) Top five KEGG terms by adjusted *p*-value (□≤0.05). (right panel) top 3 TF families (abundance) for each cluster.

There were 6,042 DEGs (adjusted *p*-value < 0.05) between R1 and R25 treatments compared to the Mock inoculation (**Data S1**). Among them, 2,387 (39.5%) overlapped between R1 and R25, whereas 175 (2.9%) and 3,480 (57.6%) DEGs were present only in R1 or R25 treatments, respectively (**Fig 2b**). To validate and contextualize our findings, we isolated total RNA from R1-, R25-, and Mock-inoculated hypocotyls (independent experiments from those used for RNA-seq), generated adapter-ligated cDNA libraries, and performed RNA hybridization-based enrichment followed by high-throughput sequencing (i.e., Capture-seq). The RNA hybridization was performed with biotinylated RNA baits designed for seven genes that showed elevated expression upon *P. sojae* infection (i.e., pathogen-induced genes) across all NILs reported in Lin et al. [27] as well as both interaction types in our RNA-seq dataset. Baits were incorporated for the six aforementioned reference genes as an internal standard. We recovered 100% of our capture library. Further, 7/7 pathogen-induced genes showed significantly elevated expression in inoculated vs Mock samples, while 5/6 reference genes showed stable expression across treatments (**Fig S1a,b**).

### Identification of defense-related TF families

Following transcriptome validation by Capture-seq, we used PlantTFDB [54] to identify TF-annotated genes in our DEG set (genes differentially expressed in at least one interaction). We found 447 (7.3% of DEGs) distributed across 43 TF families. Among these families, WRKY was the most represented by total abundance (*n* = 65 encoding DEGs), followed by ERF (*n* = 56), bHLH (*n* = 44), MYB (*n* = 37), and C2H2 (*n* = 31) (**Fig 2c**). The RAV TF family was most represented by the percentage of genome-wide proportion (60%) (**Fig 2c**). Moreover, proportions of WRKY, ERF, HSF, CAMTA, and RAV TF-encoding genes were significantly enriched in the DEG set (hypergeometric *p*-value < 0.05).

To predict functions of the defined TF families in the present pathosystem, K-Means clustering was used to segregate DEGs into nine co-expression clusters. In clusters 3 and 6 (hereafter “down-regulated clusters”), both compatible and incompatible interactions showed reduced expression in comparison to Mock, with mean expression higher in the incompatible interaction than in the compatible (**Fig 2d**). A reciprocal pattern was observed in clusters 1, 2, 4, 5, 7, 8, and 9 (hereafter “up-regulated clusters”), wherein the majority of genes were up-regulated compared to the Mock, and the compatible interaction displayed the highest mean expression (**Fig 2d**). We explored these trends by assigning functional annotations to each gene cluster with GO term and Kyoto Encyclopedia of Genes and Genomes (KEGG) pathway enrichment analyses [55,56]. Interestingly, up-regulated clusters displayed enrichment for secondary metabolism and signaling-related KEGG terms, particularly those inferring MAPK signal transduction (DEGs/Total genes: 57/229), phenylpropanoid biosynthesis (48/214), flavonoid biosynthesis (23/67), and plant-pathogen interaction (64/280) (**Fig 2e**; **Data S1**). The majority of genes corresponding to these terms were differentially expressed in both compatible and incompatible interactions compared to Mock, with the exception of plant-pathogen interaction and MAPK-related pathways, where 27 and 35 genes, respectively, were differentially expressed exclusively during the compatible interaction. Down-regulated clusters were enriched with GO and KEGG terms related to primary metabolism (e.g., photosynthesis) (**Fig 2e**; **Data S1**), which likely reflected a reallocation of cellular energy to defense [57].

Next, we examined TF representation in the co-expression clusters involved in the signaling and secondary metabolic responses to *P. sojae* (i.e., up-regulated clusters). WRKY was the most abundant TF family for four clusters (4, 7, 8, and 9), ERF for two clusters (1 and 5), and C2H2 for cluster 2 (**Fig 2e**). Furthermore, four TF families were represented in more than half of the up-regulated clusters, with ERF present in 7/7, WRKY in 6/7, C2H2 in 4/7, and MYB in 4/7.

### Comprehensive identification of TFBS

Transcriptome analysis prompted the identification of TFBS (and thereby target genes) for defense-related TF families. To this end, we performed DAP-seq for GmWRKY30, whose corresponding gene (*Glyma.06G125600*) was differentially expressed by RNA-seq. In addition, *GmWRKY30* homologs promoted resistance to hemibiotrophic and necrotrophic fungi in rice (*Oryza sativa*) [58] and to *Cucumber mosaic virus* in Arabidopsis [59]. Treatments were prepared for DAP-seq by inoculating soybean hypocotyls with an R1 *P. sojae* isolate (sample hereafter referred to as “WRKY30_P1”) or sterile media (hereafter “WRKY30_M1”) (see “Materials and Methods” for full details). WRKY30_P1 and WRKY30_M1 displayed 6,415 and 2,083 peak regions, respectively, corresponding to various genomic features (**Fig 3**; **Data S3**; **Table S2**). Motif enrichment analysis was then performed with MEME-ChIP [60] and demonstrated that bound regions present in both samples were statistically enriched for the WRKY TF binding site, W-box (TTTGAC/T), implicating that these regions were indeed bound by a WRKY TF. To establish regulatory roles of GmWRKY30 during *P. sojae* infection, we obtained the peaks annotated as promoters (defined as 1,000 bp up- and downstream of the transcription start site [TSS]) and retained the regions shared by WRKY30_P1 and WRKY30_M1 (235 promoters) as well as those found exclusively in WRKY30_P1 (1,110 promoters). Of these, 212 promoters corresponded to genes in our DEG set. Interestingly, 174/212 were present exclusively in the WRKY30_P1 sample (**Data S3**). Given that the only difference between WRKY30_P1 and WRKY30_M1 samples was DNA methylation marks, this observation suggested that the soybean genome undergoes differential methylation during *P. sojae* infection, perhaps as a mechanism to prevent autoimmunity [16]. Moreover, 35/212 target genes were annotated as TFs, indicating putative auto- and cross-regulatory activity of GmWRKY30 during the immune response. Thirteen of the 35 TF-encoding DEGs belonged to the WRKY TF family, and all were present in the up-regulated clusters (**Data S1**; **Data S3**). Furthermore, KEGG functional annotation revealed that eight GmWRKY30 targets were part of the MAPK or plant-pathogen interaction pathways described above (**Data S1**; **Data S3**). These data concomitantly suggest that GmWRKY30 regulates the expression of other TFs and signaling components during soybean-*P. sojae* interaction.

**Fig 3.**
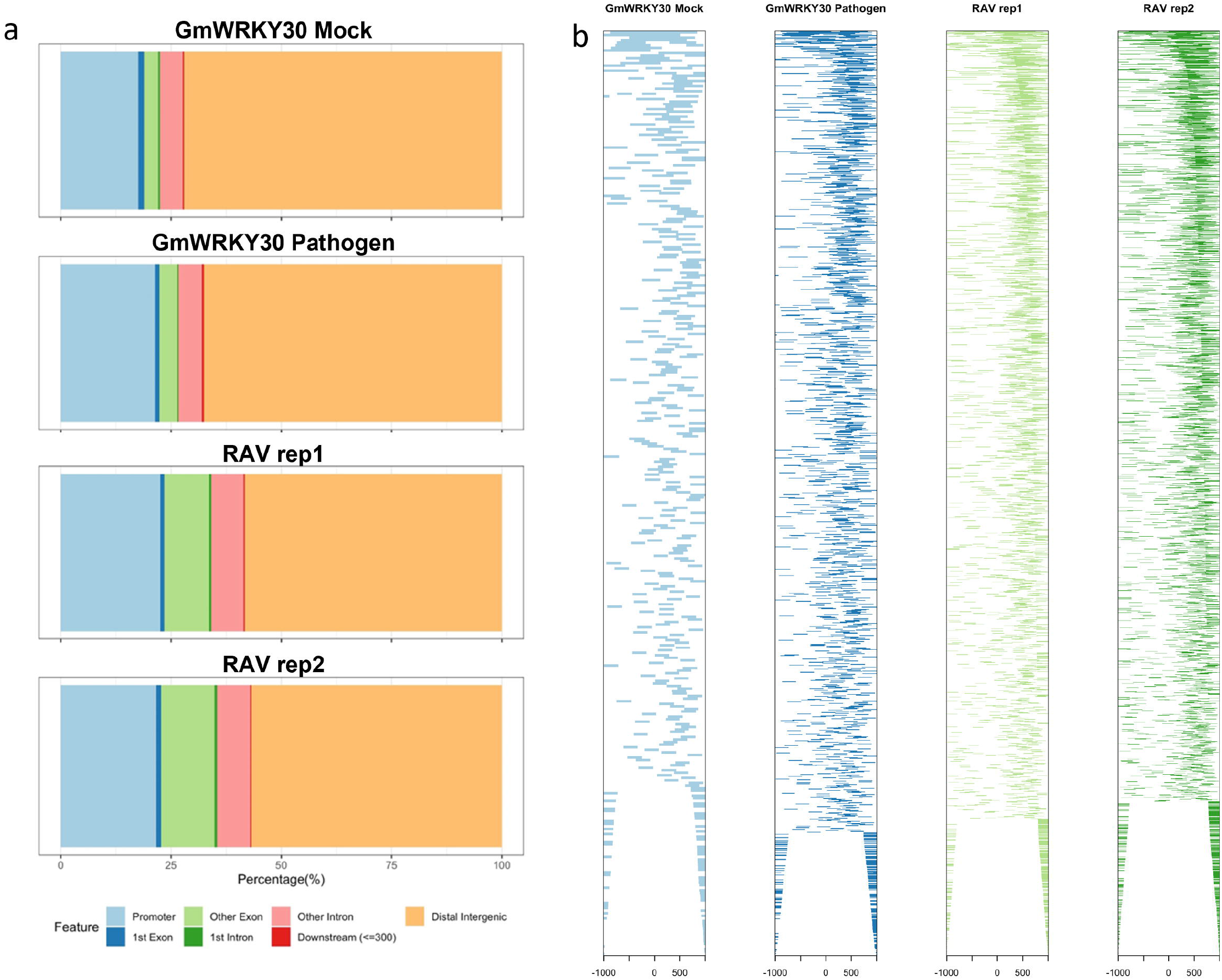
DAP-seq identification of GmWRKY30 and GmRAV TFBS. (**a**) Distribution of DAP-seq peaks across genomic features. (**b**) Distance of peaks from the TSS.

RAV was the most abundant TF family by a percentage of genome-wide proportion in our DEG set; therefore, we obtained DAP-seq data for a GmRAV TF from Wang et al. [61]. The corresponding gene (*Glyma.10G204400*) was significantly up-regulated in both compatible and incompatible interactions compared to Mock and displayed similar expression dynamics in Lin et al. [27] upon *P. sojae* infection. In the present analysis, GmRAV was bound to 3,409 promoters corresponding to 389 genes in our DEG set (**Data S1**; **Data S4**). Of these, 29 encoded TFs. One hundred seventy-six of the 389 targets (45%) were present in either cluster 3 or 6 (down-regulated clusters) (**Data S1**; **Data S4**). Cooperatively, functional enrichment demonstrated that GmRAV targets included genes relevant to photosynthesis and carbon metabolism (**Data S1**; **Data S4**), indicating that GmRAV may repress primary metabolic activity during pathogen infection.

### DNN prediction of TFBS

While TFs in a structural family have the capacity to function distinctly *in vivo*, they often share intrinsic CRE preference [62,63,64]. Therefore, we hypothesized that binding sites obtained for a single TF could be used to predict binding sites for other members of the same family. To test this hypothesis, we trained Convolutional Recurrent Neural Networks (CRNNs), which couple CNN and bi-directional long short-term memory layer architecture [65] (**Fig 4**), using peak summits of WRKY30_P1 and WRKY30_M1 samples with either 32- or 201-bp peak regions (**Fig S2**). The CRNN with a 201-bp region outperformed the CRNN with a 32-bp peak region and displayed an 89% validation accuracy, 90% test accuracy, and a false positive rate of less than 3% (**Table 1**). We trained a similar model for GmRAV, which had an 89% validation accuracy, 89% test accuracy, and a less than 3.5% false positive rate (**Table 1**). Moreover, for both GmWRKY30 and GmRAV CRNN models, the area under the receiver operating characteristic (auROC) curve was beyond 0.88, and the area under the precision-recall curve (auPRC) beyond 0.81 (**Fig S3**). To determine if a CRNN model trained for one TF could generalize to members of the same family intraspecifically, we generated AmpDAP-seq data for GmWRKY2 (homologous to AtWRKY2 and encoded by *Glyma.06G320700*). For this sample, we first observed peak distribution across genomic features (**Fig S4**; **Data S5**) and used MEME-ChIP to verify statistical enrichment of the W-box CRE within peak regions. We then used peak regions to test if the GmWRKY30 CRNN could predict GmWRKY2-bound sites. The prediction accuracy was above 82%, with a false positive rate of less than 6% (**Fig 5a**). Furthermore, we explored the interspecific generalization capacity of the GmWRKY30 CRNN by performing a cross-species prediction on AtWRKY30 (encoded by *AT5G24110;* homologous to GmWRKY30) DAP-seq data (generated by O’Malley et al. [66] and reanalyzed by Song et al. [67]). For this analysis, the prediction accuracy was above 84% with a false positive rate of less than 7% (**Fig 5a**).

**Fig 4.**
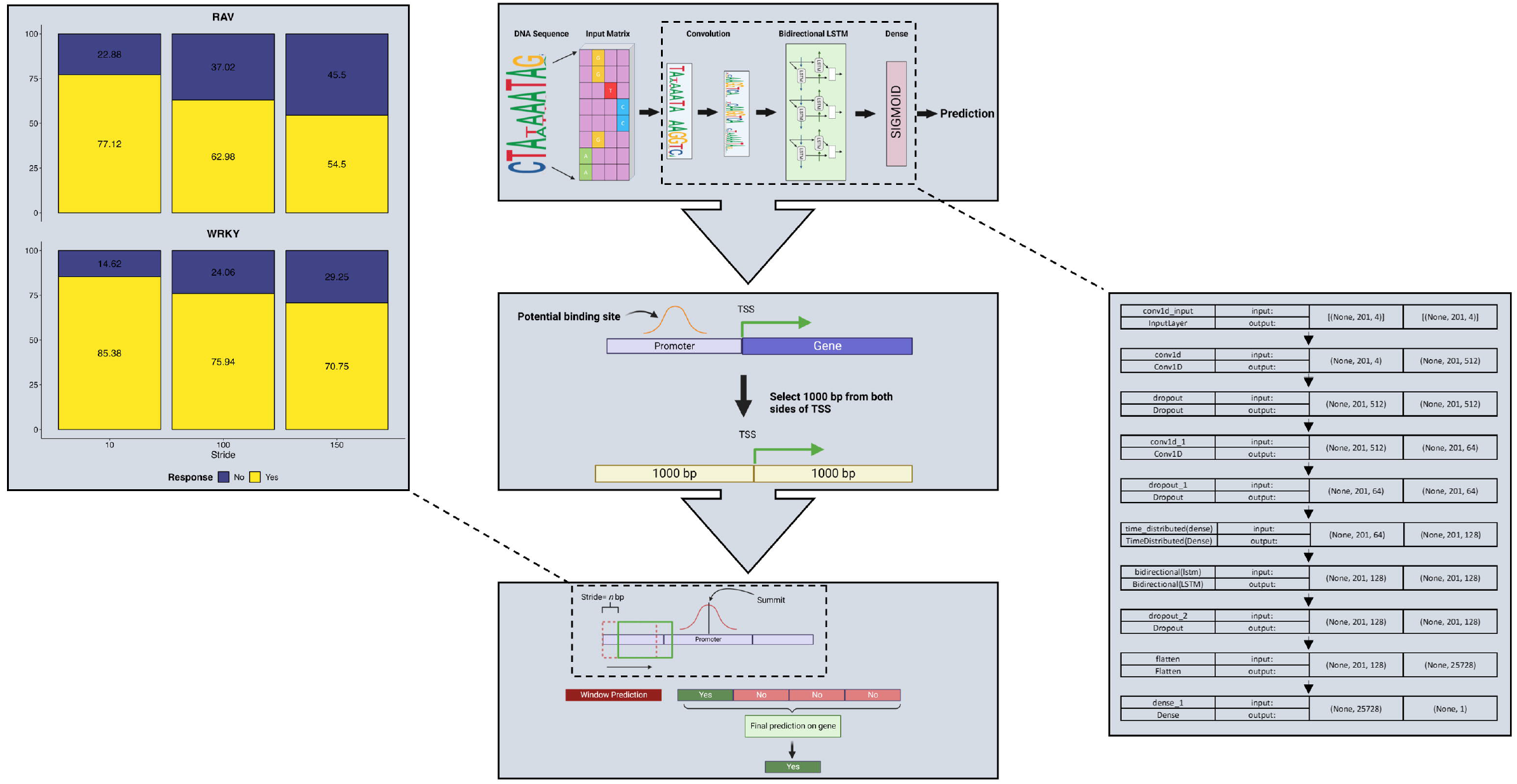
Schematic illustration of CRNN architecture and TFBS prediction.

**Fig 5.**
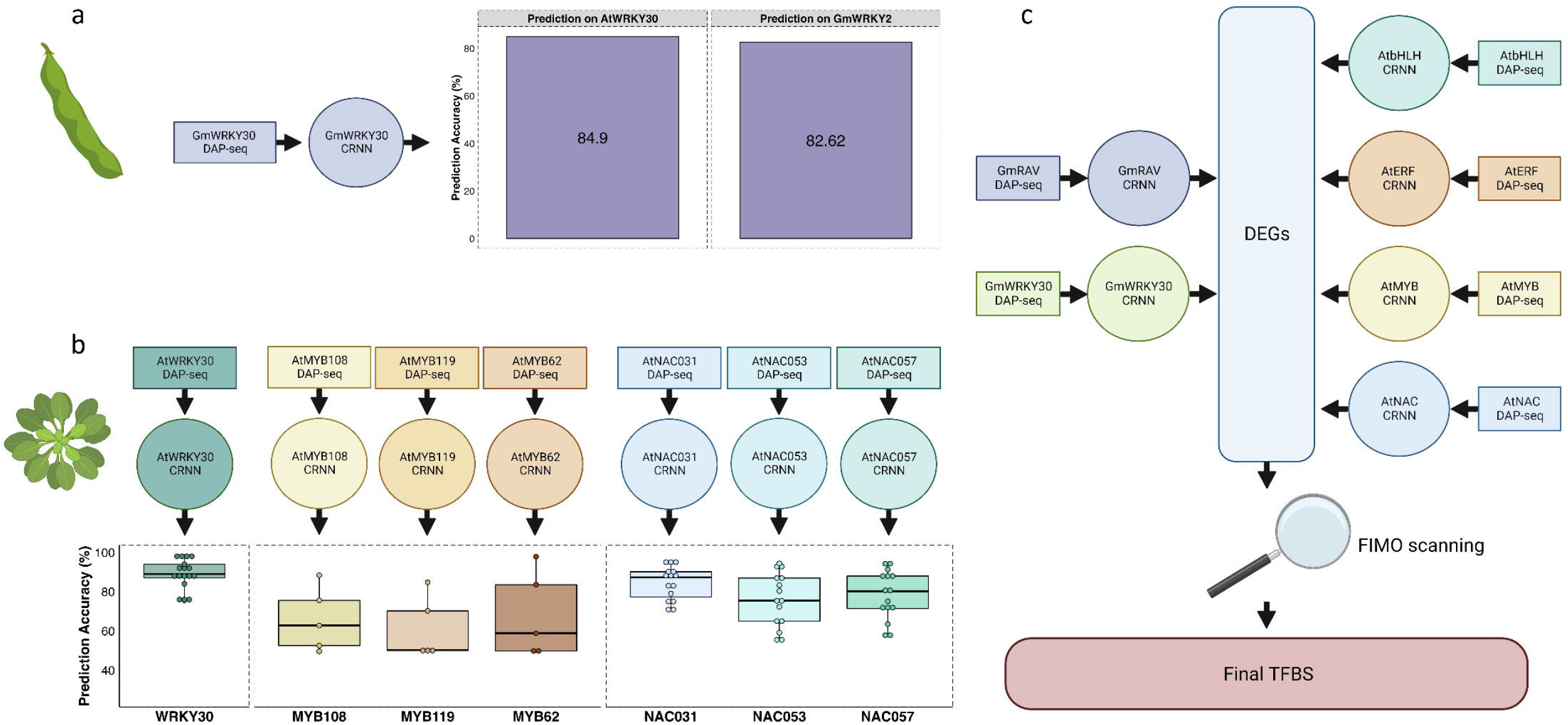
Generalization testing for soybean and Arabidopsis CRNNs and schematic illustration of TFBS prediction for defense-related TF families. (**a**) The GmWRKY30 CRNN was used to predict TFBS interspecifically with AtWRKY30 DAP-seq data (left barplot) and intraspecifically with GmWRKY2 AmpDAP-seq data (right barplot). (**b**) AtWRKY, AtMYB, and AtNAC CRNNs were trained with available DAP-seq data and used to predict binding sites for other members of their respective families. (**c**) The Arabidopsis-based models, along with the GmWRKY30 and GmRAV models, were used to predict TFBS in our DEG set. These predictions were overlaid with FIMO scans to elucidate TF-target gene interactions.

**Table 1.** Summary of GmWRKY and GmRAV CRNN models. TPR: True Positive Rate; TNR: True Negative Rate; FPR: False Positive Rate; FNR: False Negative Rate

| Model | Train Acc. (%) | Val Acc. (%) | Test Acc. (%) | TPR (%) | TNR (%) | FPR (%) | FNR (%) |
| --- | --- | --- | --- | --- | --- | --- | --- |
| **WRKY** | 90.97 | 89.77 | 90 | 33.67 | 56.21 | 2.71 | 7.41 |
| **RAV** | 89.09 | 89.2 | 89 | 44.88 | 45.33 | 3.49 | 6.3 |

Yet, it remained unclear whether these patterns would recur within/across TF families. Therefore, we utilized DAP-seq data for AtWRKY30, AtMYB62, AtMYB108, AtMYB119, AtNAC031, AtNAC053, and AtNAC057 [66,67] to train additional CRNNs. The AtWRKY30 model validation accuracy was 97%, test accuracy 97.33%, and false positive rate 0.13% (**Table 2**). The model was used subsequently to predict binding sites for 17 other AtWRKY TFs with available DAP-seq data [66,67] and presented a mean prediction accuracy of 89% with a mean false positive rate under 1% (**Fig 5b**; **Data S6**). The three AtMYB models had 92-98% validation accuracies, 91-98% test accuracies, and 0.8-2.76 % false positive rates (**Table 2**) and were used to predict binding sites for 5 AtMYB TFs [66,67], presenting a mean prediction accuracy of 64.92% and a mean false positive rate of 1.53% (**Fig 5b**; **Data S6**). Similarly, the three AtNAC models posed validation accuracies between 95.5-98%, test accuracies between 95-98%, and a false positive rate ranging between 0.6-1.79% (**Table 2**). These models were used for predicting binding sites for 15 other AtNAC TFs [66,67] with a mean prediction accuracy of 79.6% and a mean false positive rate of 1.09% (**Fig 5b**; **Data S6**). These findings indicated that a model trained using the TFBS of one family member could predict the TFBS of another member with reasonable accuracy. Therefore, GmWRKY30 and GmRAV models were used to predict TFBS on promoters of the DEGs from the transcriptome analysis (**Fig 5c**). To further reduce false positives, we scanned the same promoter regions using Find Individual Motif Occurrences (FIMO) [68] with motifs obtained from the JASPAR database [69] to find CREs. The results from the FIMO scan were overlapped with our predicted sites to get a highly confident set of TFBS. From this, we obtained 3,298 GmWRKY targets with 267 corresponding to TF-encoding genes (60% of TFs in the DEG list). Similarly, GmRAV-predicted targets included 1,925 genes, 121 of which encoded TFs (27% of TFs in the DEG list) (**Data S1**).

**Table 2.**
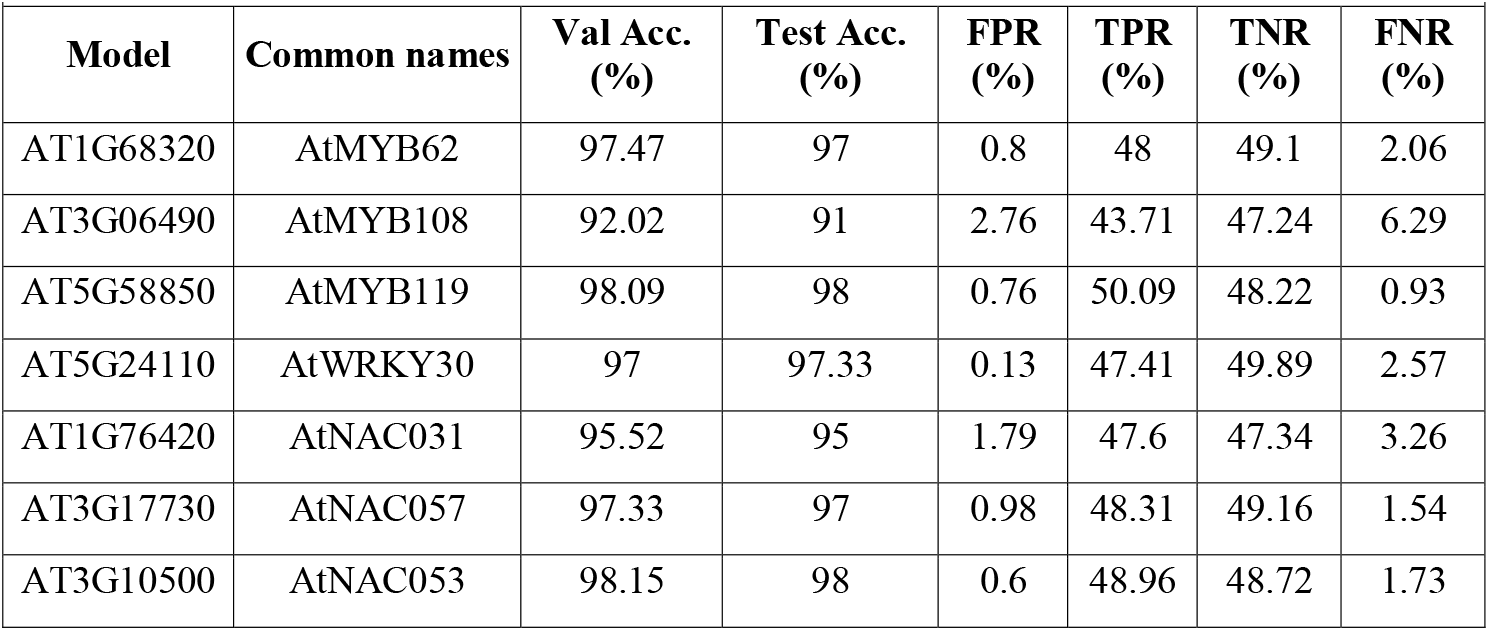
Summary of Arabidopsis data-based CRNN models used for generalization tests. TPR: True Positive Rate; TNR: True Negative Rate; FPR: False Positive Rate; FNR: False Negative Rate

| Model | Common names | Val Acc. (%) | Test Acc. (%) | FPR (%) | TPR (%) | TNR (%) | FNR (%) |
| --- | --- | --- | --- | --- | --- | --- | --- |
| AT1G68320 | AtMYB62 | 97.47 | 97 | 0.8 | 48 | 49.1 | 2.06 |
| AT3G06490 | AtMYB108 | 92.02 | 91 | 2.76 | 43.71 | 47.24 | 6.29 |
| AT5G58850 | AtMYB119 | 98.09 | 98 | 0.76 | 50.09 | 48.22 | 0.93 |
| AT5G24110 | AtWRKY30 | 97 | 97.33 | 0.13 | 47.41 | 49.89 | 2.57 |
| AT1G76420 | AtNAC031 | 95.52 | 95 | 1.79 | 47.6 | 47.34 | 3.26 |
| AT3G17730 | AtNAC057 | 97.33 | 97 | 0.98 | 48.31 | 49.16 | 1.54 |
| AT3G10500 | AtNAC053 | 98.15 | 98 | 0.6 | 48.96 | 48.72 | 1.73 |

### Cross-species prediction of soybean TFBS

Based upon the interspecies generalization capacity of the GmWRKY30 model, we wanted to further leverage the homology between Arabidopsis and soybean to build CRNNs and conduct cross-species predictions for other defense-related TF families. In doing so, we built CRNNs for AtERF, AtbHLH, AtC2H2, AtRAV, AtWRKY, AtMYB, and AtNAC TF families by combining available DAP-seq data for each family. With the exception of the AtRAV and AtC2H2 models, the training, validation, and testing accuracies were above 90%, and false positive rates less than 1.2% for all models (**Table 3**). Both AtRAV and AtC2H2 models had training, validation, and testing accuracies of less than 86% with increased false negative rates and were thus not used for subsequent analyses (**Table 3**). Next, we generated AmpDAP-seq data for GmMYB61 (encoded by *Glyma.10G142200*) (**Fig S4**; **Data S5**) and used AtWRKY and AtMYB CRNNs to perform cross-species predictions on GmWRKY30 DAP- and GmMYB61 AmpDAP-seq data, respectively. Both predictions demonstrated modest accuracies (approximately 61%), yet less than 1% false positive rates (**Table 4**). We posited that, while Arabidopsis-to-soybean predictions would likely miss some true TFBS, we could have confidence in those classified as bound. Therefore, we used AtMYB, AtERF, AtNAC, and AtbHLH CRNNs to predict TFBS for promoter regions of our DEGs (Fig 5c). Consistent with soybean CRNNs, all of the predicted TFBS were overlapped with FIMO predictions to get a highly confident set of targets.

**Table 3.**
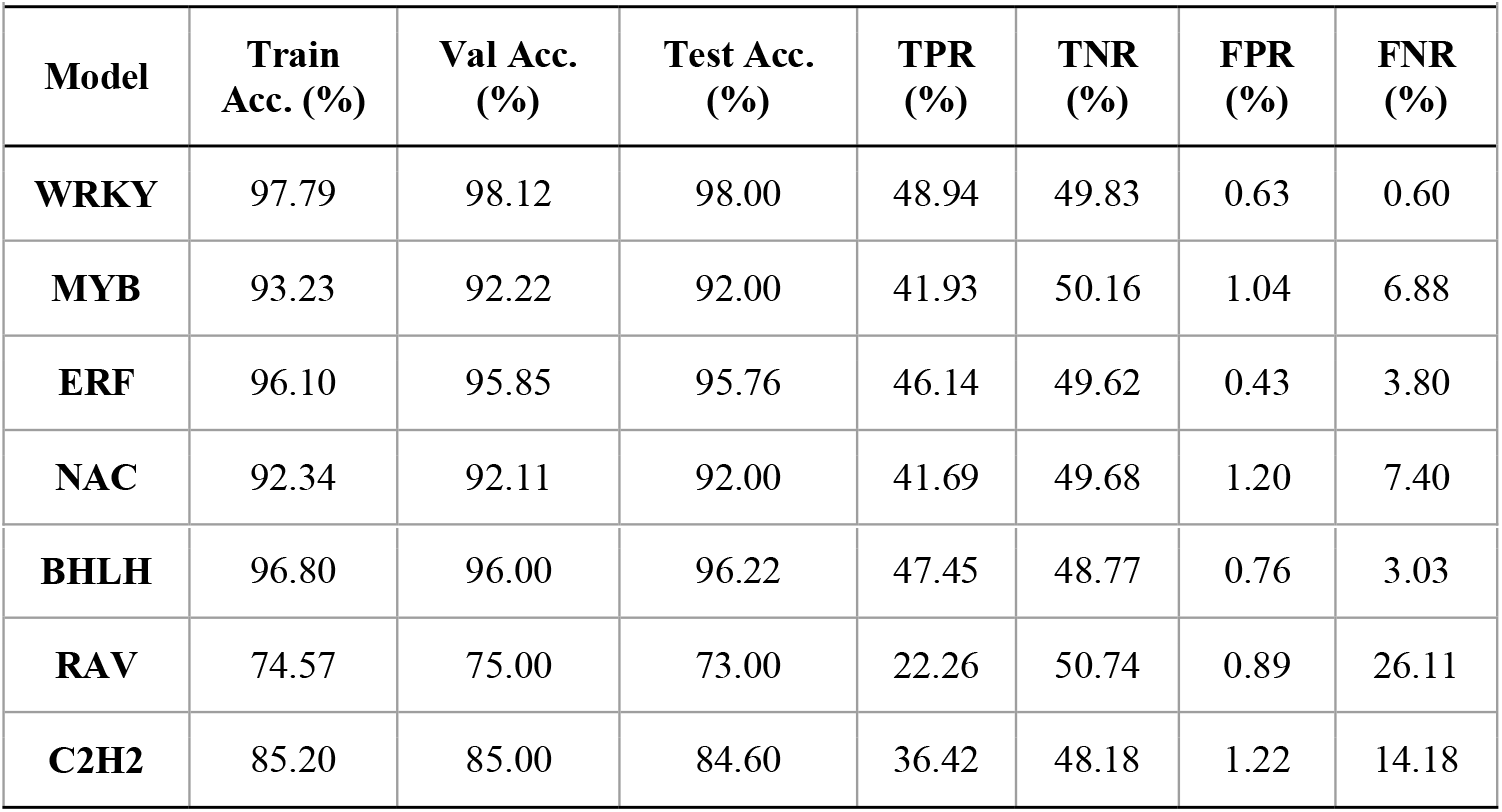
Summary of Arabidopsis data-based CRNN models used for cross-species predictions. TPR: True Positive Rate; TNR: True Negative Rate; FPR: False Positive Rate; FNR: False Negative Rate

| **Model** | **Train Acc. (%)** | **Val Acc. (%)** | **Test Acc. (%)** | **TPR (%)** | **TNR (%)** | **FPR (%)** | **FNR (%)** |
| --- | --- | --- | --- | --- | --- | --- | --- |
| **WRKY** | 97.79 | 98.12 | 98.00 | 48.94 | 49.83 | 0.63 | 0.60 |
| **MYB** | 93.23 | 92.22 | 92.00 | 41.93 | 50.16 | 1.04 | 6.88 |
| **ERF** | 96.10 | 95.85 | 95.76 | 46.14 | 49.62 | 0.43 | 3.80 |
| **NAC** | 92.34 | 92.11 | 92.00 | 41.69 | 49.68 | 1.20 | 7.40 |
| **BHLH** | 96.80 | 96.00 | 96.22 | 47.45 | 48.77 | 0.76 | 3.03 |
| **RAV** | 74.57 | 75.00 | 73.00 | 22.26 | 50.74 | 0.89 | 26.11 |
| **C2H2** | 85.20 | 85.00 | 84.60 | 36.42 | 48.18 | 1.22 | 14.18 |

**Table 4.** Arabidopsis-to-soybean cross-species prediction results. TPR: True Positive Rate; TNR: True Negative Rate; FPR: False Positive Rate; FNR: False Negative Rate

| **Table 4: Arabidopsis-to-soybean cross-species prediction results** |  |  |  |  |  |
| --- | --- | --- | --- | --- | --- |
| **Model** | **Cross Pred. Acc. (%)** | **TPR (%)** | **TNR (%)** | **FPR (%)** | **FNR (%)** |
| **WRKY** | 61.16 | 5.81 | 55.35 | 0.52 | 38.32 |
| **MYB** | 60.81 | 10.92 | 49.89 | 0.90 | 38.29 |

### GRN inference underpinning host defense

The combination of soybean and Arabidopsis CRNNs allowed the prediction of TFBS corresponding to 5,505 genes in the DEG set (**Fig 6a**). Global and family-level GRNs were thereby constructed with TFs represented by nodes and target genes by edges (**Fig 6b; Fig S5**). We then examined TF-level TFBS to prioritize nodes in the global GRN. To be considered, nodes had to possess statistically enriched binding motifs (*q*-value < 0.05) compared to a randomly shuffled input sequence (determined by the Simple Enrichment Analysis algorithm of Bailey and Grant [70]) and had to have corresponding genes expressed in the transcriptome analysis (**Fig 6d**). The 118 nodes meeting these criteria were prioritized by degree centrality, TF co-occurrence, and the expression pattern of corresponding genes. Degree centrality was determined from outdegree (number of edges directed to each node) and cumulative indegree (indegree = number of nodes to which an edge is directed; cumulative indegree = combined indegree for all edges of a node). Both measures were scale-free and displayed a power-law distribution (**Fig 6d**) as expected of GRN architecture [71]. Furthermore, TF co-operativity/co-occurrence metrics are required to effectively model causal GRNs [72]. We assessed putative TF co-occurrence with TF-COMB (Transcription Factor Co-Occurrence using Market Basket analysis) [73] and selected cosine association score as the objective similarity measure for co-occurring TF pairs (**Fig 6c**). Cumulative cosine (total cosine association score across all co-occurrences) was determined for each node. Lastly, we calculated the mean |log_2_FC| for node-corresponding genes across both interactions (R1 vs Mock and R25 vs Mock) for further prioritization. Nodes/node-corresponding genes present in the upper quartile for all four parameters were considered hubs (**Fig 6d**). A reciprocal approach prioritized edges by indegree, cumulative outdegree (combined outdegree of all nodes to which an edge is directed), sum cumulative cosine (combined cumulative cosine of all nodes to which an edge is directed), and mean |log_2_FC| across both interactions (**Fig S6**).

**Fig 6.**
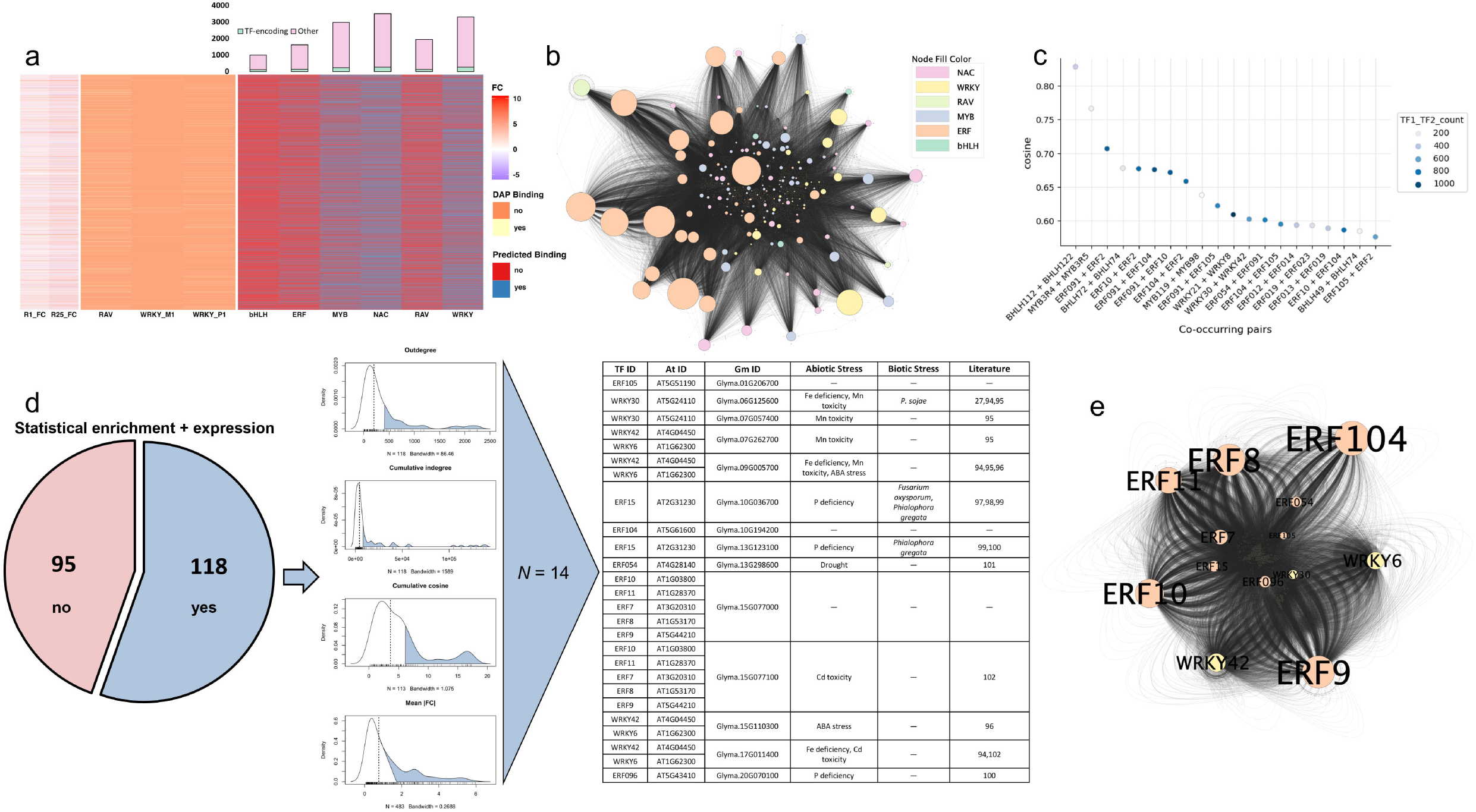
GRN inference at 24 hpi. (**a**) (left) log_2_FC (FC) of DEGs across interaction types, (middle) WRKY and RAV binding site representation in the DEG set derived from DAP-seq, and (right) binding site representation for each TF family in the DEG set derived from CRNN + FIMO prediction. The bar plot shows the total number of target genes for each family, as well as the number of TF-encoding target genes (light green). (**b**) Hairball of the global GRN. Nodes and edges represent TFs and target genes, respectively. Node size corresponds to outdegree. (**c**) Scatterplot of the top co-occurring TF pairs by cosine association score identified with TF-COMB. The datapoint color reflects the total number of shared targets for a given TF pair. (**d**) Prioritization of nodes. Nodes that were statistically enriched by Simple Enrichment Analysis and were represented in the transcriptome analysis (n = 118) were prioritized by outdegree, cumulative indegree, cumulative cosine, and mean |log_2_FC| (Mean |FC|). Blue polygons represent the upper quartile for each parameter. Fourteen genes/13 TFs were in the upper quarter for all four parameters. (**e**) Hairball of the hub nodes. Node size corresponds to outdegree.

Hub nodes corresponded to 14 genes encoding 13 TFs, all of which belonged to ERF and WRKY families (**Fig 6d,e**). Thirteen of these were differentially expressed in the transcriptome analysis and were present in co-expression clusters 1, 4, 5, and 8 (up-regulated clusters with defense-related functional annotations). Furthermore, we assessed KEGG annotations for putative hub node targets (*n* = 1,664), and 170 targets were annotated to one or more functions. Interestingly, 67% (114/170) of the targets possessed primary metabolic terms whereas 33% (56/170) possessed terms related to defense/secondary metabolism. It is therefore reasonable to hypothesize that these hub nodes are core components of the defense-growth tradeoff in soybean by regulating transcriptional reprogramming.

## Discussion

*P. sojae* is a yield-devastating soybean pathogen subject to rapid genetic diversification and expansion within and across production environments. To unravel regulatory signatures of host defense during *P. sojae* infection, we coupled multi-omic and computational analyses to identify TF-target gene interactions at 24 hpi. In doing so, we conducted the first comparative transcriptomic study for compatible and incompatible soybean-*P. sojae* interactions within a single host genotype. Similar gene expression profiles were observed across the interaction types, implicating significant overlap between PTI- and PTI + ETI-mediated defense at 24 hpi. Enkerli et al. [74] revealed ultrastructural differences between compatible and incompatible soybean-*P. sojae* interactions at 4 hpi, with programmed cell death and impedance of hyphal growth evident in the incompatible interaction by 15 hpi. Thus, it is likely that maximal expression of PTI-potentiating transcripts during incompatibility precludes or at least precedes the *P. sojae* transition from biotrophy to necrotrophy (12-24 hpi [75,76]), and succedent activity reflects a reduction in hypersensitivity required to offset fitness costs of induced resistance. This concept draws parallels to other pathosystems in which compatible and incompatible interactions displayed consistent trends in gene expression, with the incompatible interaction eliciting a heightened, more immediate immune response [77,78,79]. A complementary explanation is that PTI/ETI convergence is attributed to the *P. sojae* arsenal present during both compatible and incompatible interactions, including MAMPs and a conserved suite of effectors [80]. Moreover, overlapping expression profiles may correspond primarily to PTI signatures not targeted by *Avr*-encoded effectors. Nevertheless, findings herein necessitate the investigation of compatible and incompatible interactions in tandem to elucidate *Rps* gene-exclusive defense mechanisms.

Plant immune signaling is remarkably tunable yet robust, allowing the co-ordination of defense and growth in a manner that maximizes host fitness [21,81,82]. Prior studies suggest signal integration underpinning the defense-growth trade-off is imposed by TF regulatory networks that modulate immune responses through transcriptional reprogramming [17]. In the present study, K-means clustering of DEGs rendered nine gene co-expression clusters, seven of which were up-regulated and corresponded to defense-related functional annotations (e.g., MAPK signaling and secondary metabolism). Cooperatively, these clusters were enriched with statistically overrepresented TF families (i.e., MYB, WRKY, NAC, ERF, and C2H2) known to regulate plant secondary metabolism [22,23] and reported in prior soybean-*P. sojae* studies [29,30,32,33,34,35,37,83]. Furthermore, WRKY and ERF were the most abundant TF families across the seven clusters. This is consistent with findings in Arabidopsis where MAPK-WRKY and MAPK-ERF complexes regulated core immune signaling through transcriptional reprogramming [17,20,84]. The remaining gene co-expression clusters were down-regulated and demarcated by growth and reproductive functional terms. Interestingly, DAP-seq suggested that gene targets of RAV (the most represented TF family in the DEG set by a percentage of genome-wide proportion) were enriched in these two clusters. This finding is consistent with prior studies wherein GmRAV has been reported to play a role in photosynthesis, senescence, abiotic stress tolerance, and phytohormone-mediated signaling [85,86,87] and act as a transcriptional repressor to delay flowering [61]. To our knowledge, the present study is the first to propose a function for GmRAV during immunity, where it acts as a repressor of primary metabolism. Thus, our transcriptome analysis evidences transcriptional reprogramming governing the defense-growth trade-off in soybean upon *P. sojae* infection.

DAP-seq data were generated/obtained for the most represented TF families by total abundance and percentage of genome-wide proportion in the transcriptome analysis, and promoter-localized DAP-seq peaks were used to train CRNNs (DNNs composed of convolutional and recurrent layers) for the prediction of novel TFBS. We leveraged CRNNs for their capacity to learn randomly composite, predictive sequence patterns [47]. Previous studies suggest that binding site motifs, along with nearby sequence features and their organization in the genome, play a vital role in TF binding. The convolutional filters in CRNNs can capture and train these binding site motifs and nearby sequence features, while recurrent layers can learn their multidimensional organization [47]. In addition, such hybrid model architecture has been used successfully to predict TF-target interactions with human data [65,88,89]. Here, our CRNN models were capable of predicting TFBS for the selected TF families in soybean and Arabidopsis with ~90% accuracy. The exclusive use of DEG promoters for binding site prediction increased the likelihood that targets were biologically valid, as the correlation between stable TF binding and TF regulation is vastly inconsistent and oftentimes poor [72,90,91]. Moreover, CRNNs trained for one TF could find TFBS for other members of the same family. We supported this notion in soybean by generating binding site data for a second WRKY TF and using the pre-existing WRKY CRNN to predict its targets. We validated our findings in another species by training additional CRNNs with Arabidopsis DAP-seq data for WRKY, MYB, and NAC TFs and predicting TFBS for various members of the respective families. In every instance, CRNNs were capable of generalizing with acceptable accuracy, posing significant potential as an alternative to wet lab-based TFBS assays. Altogether, findings herein reflect the ability to train highly accurate CRNNs for the prediction of TFBS in plants.

The DNA sequence preference of TFs is largely conserved across phylogenetically-related species, leading to the advent of deep learning-based approaches for cross-species TFBS prediction [47,64,91]. However, recent attempts at mouse-to-human/human-to-mouse and maize-to-soybean cross-species predictions suffered from high false positive rates [47,49]. We hypothesized that we could overcome such limitations due to the evolutionary proximity of soybean and Arabidopsis (two diploid, dicotyledonous species). In the present study, Arabidopsis-to-soybean predictions had moderate accuracy (approximately 60%) with low false positive rates (less than 1%). Interestingly, soybean-to-Arabidopsis predictions displayed a higher accuracy than the Arabidopsis-to-soybean. Nitta et al. [92] investigated TFBS conservation between *Drosophila* and mammals, finding that novel binding site specificities could arise via gene duplication and subsequent divergence. Perhaps the lower Arabidopsis-to-soybean prediction accuracies reflect the expansion of WRKY and MYB families in soybean, rendering soybean-specific CRE preferences. Furthermore, the lower prediction accuracies may indicate contributions to TF genomic occupancy beyond DNA sequence affinity (e.g., chromatin state; presence of cofactors), which have been demonstrated to significantly influence immunity-related transcriptional dynamics in plants [16]. Thus, the integration of complementary information (e.g., ATAC-seq data) into existing CRNN frameworks will allow for more accurate model training in the future.

We predicted TFBS for WRKY, RAV, NAC, ERF, bHLH, and MYB TF families with soybean- and Arabidopsis-based CRNNs, overlaid CRNN predictions with FIMO scans, and constructed global and family-level GRNs. Within a GRN, some TFs act as hubs to regulate many genes, while target genes are typically regulated by multiple TFs [72]. Therefore, we identified hub nodes, which presumably have an inordinate effect on phenotype [93], by integrating motif enrichment analysis, degree centrality, TF co-occurrence, and gene expression metrics. Interestingly, all hub-corresponding genes encoded WRKY or ERF TFs, suggesting these families are central components of the host immune response at 24 hpi. One could attribute this in part to the use of intraspecies prediction for WRKY and RAV and cross-species prediction for the other TF families, the latter of which was prone to high false negative rates that likely influenced degree centrality. Yet, WRKY and ERF were the most represented TF families in defense-related DEG clusters derived from the transcriptome analysis, which, when coupled with functional annotations of hub node targets, reinforced the pertinence of the two families for host defense. Moreover, the majority of hub genes identified here demonstrated differential expression in soybean upon various biotic and abiotic stresses (**Fig 6d**) [27,94,95,96,97,98,99,100,101,102]. Additional efforts must be used to functionally validate TF-target predictions, which remains a bottleneck in GRN research [72,103].

Nevertheless, this research poses limitations that must be considered. First, the inoculation procedure of Dorrance et al. [50] was used in the present study: hypocotyl wounding occurred prior to the placement of a mycelial slurry (sterile media for Mock tissues). It is possible that mechanical tissue disruption increased damage-associated molecular pattern (DAMP)-related signaling, and that DAMP + MAMP perception triggered PTI beyond what occurs *in situ* [104]. Second, the existing network lacks the temporal resolution required to fully elucidate dynamic phenomena such as plant-pathogen interactions [72]. Thus, future experiments will include more natural inoculation procedures, the integration of time-series expression and epigenomic data into CRNN frameworks, the evaluation of additional defense-relevant TF families, and the molecular investigation of *P. sojae* effector targets.

## Conclusions

In summary, TF families emphasized in this study (particularly ERF, WRKY, and RAV) are likely core components of sensory regulatory networks required for the balance of primary and secondary metabolic responses to *P. sojae* infection. Interactions predicted here shed light on convergent and discrete transcriptional complexes during host immunity and provide a framework for data integration, functional validation, and the prediction of novel regulatory components for disease resistance [21]. We also provide a framework for improved TFBS prediction by coupling high-throughput sequencing data and CRNNs. Consequently, the information herein may prove useful for the circumvention of *P. sojae* pathogenicity through the modulation of defense-related pathways and the resultant derivation of disease-resistant soybean genotypes.

## Materials and Methods

### Biological materials, pathogenicity testing, and RNA isolation

For all experiments, soybean and *P. sojae* materials were generated at Iowa State University in the lab of Dr. Alison Robertson. Axenic-grown *P. sojae* isolates were transferred from a 20% clarified vegetable juice (V8) medium onto a soft-diluted V8 medium and incubated at 25°C in the dark as defined by Dorrance et al. [50]. Concomitantly, seeds of soybean varieties Williams 82 (*Rps1k*) and Williams (*Rps*) were grown on moistened germination paper at 25°C, 16 h d^-1^ light, and 90% relative humidity. On day 7 of plant growth, seedling hypocotyls were incised 1 cm below the cotyledonary node with a sterile razor blade and inoculated with 0.1 mL mycelial slurry of a *P. sojae* Race 1 isolate (avirulent on *Rps1k*), a Race 25 isolate (virulent on *Rps1k*), or sterile media. At 24 hpi, the mycelial slurries were washed off with deionized water, and a 2 cm fragment of each hypocotyl was cut and immediately frozen in liquid nitrogen.

For each inoculation, 10 or more seedlings per treatment were kept 7 d to monitor disease development. Seedlings were scored as asymptomatic or symptomatic depending on the absence/presence of lesions and necrotic tissue in at least 90% of replicates. For an inoculation to be successful, Williams seedlings displayed disease symptoms when infected with either *P. sojae* pathotype. Williams 82 seedlings displayed hypersensitivity upon inoculation with Race 1 and were symptomatic upon infection with Race 25. Moreover, mock inoculations rendered asymptomatic, clean wounds for both varieties.

Total RNA was isolated from frozen Williams 82 hypocotyls with the NEB Monarch Total RNA Miniprep Kit (Cat #T2010S) and quantified using a Qubit fluorometer paired with the RNA high-sensitivity assay kit (Cat #Q32852). RNA purity was estimated from A260/A280 and A260/A230 ratios using a NanoDrop ND-1000 spectrophotometer (Thermo Fisher Scientific) and further assessed by gel electrophoresis (1% agarose gel at 120V for 40 min). Samples were stored at −80 °C until use.

### RNA-seq

Total RNA was sent to Novogene Corporation (Sacramento, CA, USA) for library preparation and sequencing. RNA purity was assessed using a NanoPhotometer^®^ spectrophotometer (Implen, Westlake Village, CA, USA). RNA integrity and quantitation were monitored with the RNA Nano 6000 Assay Kit (Cat #5067-1511) of the Agilent Bioanalyzer 2100 system (Agilent Technologies, CA, USA). Following quality control, cDNA libraries were prepared from 1 μg total RNA using the NEBNext Ultra™ II RNA Library Prep Kit for Illumina (Cat #E7770S) paired with the NEBNext Poly(A) mRNA Magnetic Isolation Module (Cat #E7490) following manufacturer’s instructions. Library quality was assessed on the Agilent Bioanalyzer 2100 system, and libraries were sequenced on an Illumina platform.

Resultant short reads were processed using fastp software (v0.20.1) [105] for the removal of adapter sequences and low-quality reads (Phred <33). Clean reads were then mapped to the soybean reference genome (Gmax_508_v4.0.softmasked) with HISAT2 (v2.0.5) [106]. FeatureCounts (v1.5.0-p3) [107] was used to summarize read counts per gene. Only the genes that had a mean count > 20 across all samples were considered for further analyses to improve the sensitivity for differential gene expression analysis. The batch-level bias was removed using Combat-Seq in the Bioconductor R package sva (v3.44.0) [51], and differential gene expression analysis was performed using the DESeq2 package (v1.34.0) [108]. Genes with expression significantly changed in the R1 and R25 treatments compared to the Mock treatment (adjusted *p*-value □≤0.05) were deemed DEGs. Furthermore, we used two experimental batches as a factor for the DESeq2 model, and any genes with an adjusted p-value □≤0.05 between the two batches were removed from further analyses.

The 6,042 genes in our DEG set were clustered using the Bioconductor R package coseq (v1.17.2) [109,110]. The batch effect-corrected count data were used as the input for the coseqR and centered log-ratio-transformation and trimmed means of M values normalization were performed to normalize the counts. Genes assigned to each cluster were used for visualization and GO and KEGG enrichment analyses. For GO enrichment analysis, GO terms were downloaded from the Gene Ontology Meta Annotator for Plants (GOMAP) database [111], and the enrichment analysis was performed for each co-expression cluster using the TopGO Bioconductor package (v2.48.0) [112]. GO terms with a *p*-value □≤0.05 were considered enriched terms. Similarly, for the KEGG enrichment analysis, terms were downloaded from the KEGG database, and enrichment analysis was performed using the clusterProfiler Bioconductor package (v4.4.4) [113]. KEGG pathways with an adjusted *p*-value □≤0.05 were considered overrepresented. Genes encoding putative TFs were annotated using PlantTFDB [54]. To calculate the statistical significance between observed TF abundance in our DEG set and their genome-wide proportions, we conducted a proportion test using the prop.test function in RStudio (v3.6.3) [114]. Data visualization was performed using ggplot2 (v3.3.6) [115].

### Capture-seq

Capture-seq was used to validate the expression of pathogen-induced genes and internal standards (*n* = 13 genes). Hypocotyl inoculation and RNA isolation were performed as described above. Adapter-ligated cDNA libraries were then prepared from total RNA using the NEBNext Ultra™ II RNA Library Prep Kit for Illumina (Cat #E7770S) following appendix modifications for size selection of 300 nt inserts (420 nt final library size). In doing so, mRNA was fragmented using First-Strand Synthesis Reaction Buffer and Random Primer Mix (2X) at 94°C for 10 min (compared to 15 min in the protocol). Moreover, the incubation time during first-strand cDNA synthesis was increased from 15 to 50 min at 42°C. Size selection of libraries was performed using 25 and then 10 μl of Agencourt AMPure XP beads (Beckman Coulter, Brea, CA, USA, Cat #A63880). All libraries were quantified with a Qubit fluorometer using the dsDNA high-sensitivity assay kit (Cat #Q32851) and visualized by gel electrophoresis (2% agarose gel at 100V for 60 min).

120-nt biotinylated RNA baits were designed by Integrated DNA Technologies (IDT, Coralville, IA, USA) and encoded the sense strand of the genes of interest. Sequence capture was then performed as described in the xGen hybridization protocol from IDT (http://sfvideo.blob.core.windows.net/sitefinity/docs/default-source/protocol/xgen-hybridization-capture-of-dna-libraries.pdf?sfvrsn=ab880a07_6) with modifications. 500 ng of each barcoded Illumina library was pooled into a single 1.5 mL low-bind microcentrifuge tube and combined with 5 μg of salmon sperm DNA (Cat #15632011) and 2 μl of xGen Universal BlockersTS Mix (Cat #1075474) to prevent bait hybridization with repetitive elements/adapter sequences. Samples were dried for ~90 min in an ISS110 SpeedVac System (Thermo Fisher Scientific). Pelleted material was resuspended in 8.5 μl of 2X hybridization buffer, 2.7 μl of Hybridization Buffer Enhancer, 4 μl of a working bait stock (100 attomoles/bait/μl), and 1.8 μl of nuclease-free water to a final volume of 17 μl. A hybridization reaction was then performed with an initial denaturation at 95°C for 30 s followed by a 16-hr incubation at 65°C. M-270 Streptavidin beads were equilibrated, pelleted by a magnet, and mixed with the hybridization product for another 45-min incubation at 65°C. Heated and room temperature washes were performed as recommended and 20 μl of nuclease-free water was added to the beads. Post-capture PCR was then performed by adding the following components to the capture product: 1.25 μl of xGen library amplification primer, 10 μl of 5X Phusion HF Buffer, 1 μl of 10 mM dNTP, 0.5 μl Phusion High-Fidelity DNA Polymerase (Cat #M0530S), and 17.25 μl of nuclease-free water to a final volume of 50 μl. Amplification settings included polymerase activation at 98°C for 45 s followed by 10 cycles of denaturation at 98°C for 15 s, annealing at 60°C for 30 s, extension at 72°C for 30 s for each cycle, and a final extension at 72°C for 1 min. PCR fragments were purified using Agencourt AMPure XP beads (Cat #A63880) and eluted using 0.1X Tris-Ethylenediamine Tetraacetic Acid. Captured libraries were quantified with a Qubit fluorometer using the dsDNA high-sensitivity assay kit (Cat #Q32851). Library quality was assessed on the Agilent Bioanalyzer 2100 system (Agilent Technologies, Santa Clara, CA, USA) at Novogene Corporation. Samples were then sequenced on an Illumina platform with 150-bp paired-end reads.

The resultant short reads were preprocessed to remove poor-quality reads/sequencing artifacts using Cutadapt (3.0) [116] and BBDuk (https://sourceforge.net/projects/bbmap/) (Phred <30). The preprocessed short reads were aligned to soybean primary transcripts (Gmax_275_Wm82.a2.v1.transcript_primaryTranscriptOnly.fa) obtained from Phytozome [117] using Kallisto pseudoaligner (v0.46.1) [118] with default parameters. Differential gene expression analysis and further data processing were performed in RStudio (v3.6.3) [114] as described for the RNA-seq analysis.

### DAP- and AmpDAP-seq

DAP- and AmpDAP-seq experiments were conducted as described previously [42]. The open reading frame of each TF was cloned independently into a Gateway-compatible pIX-HALO expression vector containing an N-terminal HaloTag (Arabidopsis Biological Resource Center, stock #CD3-1742). Protein complexes were then expressed *in vitro* using the TNT SP6 Coupled Wheat Germ Extract System (Promega, Madison, WI, USA, Cat #L4130) and purified using Magne HaloTag Beads (Cat #G7282). To prepare DAP-seq samples, hypocotyls were inoculated with a mycelial slurry from a *P. sojae* Race 1 isolate or sterile media as described above. Tissues were collected at 24 hpi, and genomic DNA was isolated using the Zymo Research *Quick-DNA* Plant/Seed Miniprep kit (Cat #D6020, Irvine, CA, USA) with the addition of 2-mercaptoethanol. Three replicates per treatment were pooled to minimize biological variation, and ~5 μg DNA per pool was fragmented and ligated with modified Illumina adapters. Adapter-ligated fragments were then incubated with an immobilized HALO-tagged TF protein. For AmpDAP-seq, fragments were PCR-amplified prior to incubation with a TF complex, permitting binding in the absence of secondary modifications. In both instances, bound DNA was eluted and indexed using unique barcoded primers during PCR enrichment. Indexed libraries were quantified with a Qubit fluorometer paired with a dsDNA high-sensitivity assay kit (Cat #Q32851). Following quantification, libraries were sequenced on an Illumina platform with 150-bp paired-end reads at the University of Arkansas for Medical Sciences (Little Rock, AR, USA).

Short reads were aligned to the soybean reference genome Williams 82 Assembly 4 Annotation 1 (Gmax_508_v4.0.softmasked) using Burrows-Wheeler Alignment tool (v0.7.17-r1188) [119] and duplicated reads were removed using sambamba (v0.6.8) [120]. DAP-seq peaks were called using the Model-based Analysis of ChIP-Seq peak caller (v2.2.7.1) [121] with empty vector samples as the background control. Bioconductor packages ChIPQC (v1.8.2) [122] and ChIPseeker (v1.32.0) [123] were used to assess the DAP-seq data quality and peak annotation, respectively. The peak annotation was performed using Williams 82 Assembly 4 Annotation 1 (Gmax_508_v4.0.softmasked) genome annotation.

### Data preprocessing for CRNNs

To predict TFBS, CRNNs were trained with the aforementioned GmWRKY DAP-seq data. We combined the peak regions from both samples using pandas (v1.4.0) [124] and pybedtools (v0.9.0) [125] to obtain a non-redundant set of peak regions. Similarly, we obtained DAP-seq data for GmRAV from Wang et al. [61] and performed peak calling as mentioned above. The two biological replicates from the latter study were pooled together to get a non-redundant list of peaks. Sequences corresponding to 201-bp peak summit regions were obtained using the soybean reference genome with soft masked (Gmax_508_v4.0.softmasked obtained from Phytozome) and the same genome assembly version used for all other processing and annotations. A negative data set was created from the shuffle tool from BEDTools (v2.30.0) [126], excluding positive binding site regions equal to an exact number of positive binding sites with an identical bp length. The masked regions and low-intensity sequences were removed from the FASTA files. **Fig S2** illustrates the selection of 201-bp bound and unbound sites. The bound sites were given the label “1” (Positive), and unbound sites were labeled “0” (Negative). Both positive and negative data sets were combined to obtain the complete data set for each TF. Next, the sequences were converted into one-hot encoded binary information. In the one-hot encoding process, we consider the DNA sequence as a one-dimensional sequence represented by 4 binary channels. The encoding was conducted as follows: A = (1 0 0 0), C = (0 1 0 0), G = (0 0 10), and T= (0 0 0 1). The input for each TF was (*n*, 201, 4) three-dimensional array where *n* is the number of sequences for each data set.

For Arabidopsis, Song et al. [67] reanalyzed the DAP-seq data from O’Malley et al. [66], and we obtained the 32-bp peak summit regions from the reanalyzed dataset. Based on Akagi et al. [48], 32-bp summit peak regions perform the best for Arabidopsis DNN models. Therefore, similar to soybean, 32-bp sequences corresponding to positive and negative sets were obtained from the Arabidopsis genome sequences (genome version: TAIR 10; downloaded from The Arabidopsis Information Resource; TAIR) [127].

### Model architecture, training, and testing

We used CRNN architecture for both soybean and Arabidopsis models. The input to the CRNN network was the one-hot encoded 201-bp window of DNA sequence, which was passed through a convolutional layer with 256 20-bp filters with a Rectified Linear Unit (ReLU) activation. The next layer was a convolutional layer with 64 8-bp filters with ReLU activation. This was followed by a 128-node time-distributed layer with ReLU activation. Next was a bi-directional long short-term memory network with 64 internal nodes, followed by a 50% Dropout layer. The final layer was a single sigmoid-activated neuron. See **Fig 4** for full architecture.

For the model training, data sets were split into 70% training, 15% validation, and 15% testing. The training was conducted with Keras v2.8 [128] with backend TensorFlow (v2.8) [129] using the Adam optimizer with a 0.001 learning rate. The training ran for 100 epochs. Early stopping with a patience of 20 was used to prevent overfitting. The models were trained with a batch size of 256.

### Testing the capacity of models to generalize across TF families

We wanted to test the ability of a model trained using binding site data for one TF to predict binding sites across the TF family. First, we obtained AmpDAP-seq data for GmWRKY2 and obtained 201-bp peak summit regions and their corresponding DNA sequences as described above. Next, we used these as input for the model trained using GmWRKY30. Model accuracy was calculated based on the number of correct predictions/number of bound sites X 100.

Given the limited amount of soybean binding site data, we used Arabidopsis DAP-seq data to further verify model generalization capacity. We trained models using Arabidopsis DAP-seq data for AtWRKY30 (*AT5G24110*), AtMYB62 (*AT1G68320*), AtMYB108 (*AT5G58850*), AtMYB119 (*AT3G06490*), AtNAC031 (*AT1G76420*), AtNAC053 (*AT3G10500*), and AtNAC057 (*AT3G17730*) [66,67] using the methods described above. Similar to soybean, 201-bp peak summit regions for 17 WRKY family members, 15 NAC family members, and five MYB family members were used as input for each model. Model predictions were recorded for each family member and model accuracy was calculated as described above.

### Cross-species predictions using the Arabidopsis DAP-seq data

We built models for AtWRKY, AtMYB, AtERF, AtNAC, AtBHLH, and AtC2H2 TF families using Arabidopsis DAP-seq peaks. Akagai et al. [48] used 32-bp peak summit regions to train models with combined DAP-seq data to conduct cross-species prediction. Therefore, we adopted a similar approach and created a combined data set for each TF family to train models using 32-bp peak summit regions (with the exception of AtRAV, for which we obtained data from the ReMap database [130]). In total, we combined DAP-seq data from 24 WRKY, 19 MYB, 17 NAC, 13 ERF, 4 C2H2, and 3 bHLH to create family-level datasets. Further, we trained models using 201- and 32-bp peak regions. Both model types showed high accuracies and low false positive rates. However, since Akagai et al. [48] used 32-bp peak summit regions to train models to conduct cross-species predictions with success, we opted for this window size for our cross-species prediction. Nevertheless, our model architecture remained the same as the 201-bp models that were used for soybean DAP-seq data, while the convolution layer filter lengths were different (in the first CONV layer, filter length [Kernel] was 20 for soybean models and ten Arabidopsis models).

To test the ability of Arabidopsis models to successfully predict soybean binding targets, we used AtWRKY- and AtMYB-trained models and 201-bp summit peak sequences obtained from soybean DAP-seq (GmWRKY30) or AmpDAP data (GmMYB2) as the input for each model. Model prediction was compared with true TFBS to get cross-species prediction accuracies.

### Predicting new gene targets

For the target prediction, we selected 1,000 bp regions on both sides of the TSS (**Fig 4**), as it has been shown that binding sites can reside on either side of the TSS [67]. Next, we obtained predictions on *n* bp (201- or 32-bp) sliding windows throughout the 2-kb sequence. Then, based on widow predictions, we determined if the gene is a potential target gene for a specific TF. If at least one window had a positive prediction, we considered that a potential target.

When obtaining predictions on the target gene, the biggest challenge is finding the optimal stride length for the sliding window. For soybean models, we tested three stride lengths: 10, 100, and 150 bp. To evaluate the best stride, we obtained predictions on 201-bp sliding windows throughout the 2-kb sequence using each stride for genes with GmWRKY30 and GmRAV binding sites on the promoter regions. We then calculated the model prediction with true bindings to get the best stride (**Fig 4**). In both models, the 10-bp stride has the highest true positive counts. Therefore, we selected the 10 bp as the optimal stride with a 201-bp window region. Leveraging the optimal stride length, we predicted binding sites using GmWRKY30 and GmRAV models for all 6,042 DEGs. If at least one window had a positive prediction, we considered the gene a potential target of the TF. Similarly, we used AtMYB, AtNAC, AtERF, and AtbHLH models to predict binding sites for their respective families in soybean.

The window regions that were predicted to contain TF binding sites were scanned using FIMO software [68] from the MEME suite (v5.0.5) [131] using the default parameters. To retrieve matches, we obtained MEME core plants position frequency matrix files from the JASPAR database [69], and any motif with a *q*-value (Benjamini–Hochberg corrected *p*-value) <0.01 was used for the overlap analyses. We used pybedtools (v0.9.0) (with options: a_and_b = a.intersect(b,wb=True)) [125] to get the overlaps between FIMO-predicted and CRNN-predicted binding sites. Then, we annotated the overlapped binding sites using their Arabidopsis trans-acting factor information obtained from the JASPAR database. Global and family-level GRNs were visualized using Cytoscape (v3.9.1) [132].

### Motif enrichment analysis

Sequences corresponding to TFBS were used as input for Simple Enrichment Analysis (v5.4.1) [70] to find statistically enriched motifs with a randomly shuffled input sequence used as the background. Soybean homologs for the Arabidopsis TFs were derived from the gene families of the PANTHER classification system [133].

### TF co-occurrence

Motif co-occurrence analysis was performed with Transcription Factor Co-Occurrence using Market Basket analysis (TF-COMB) Python module [73] with default parameters (except for the count_within() function where default options were changed to max_dist=50, binarize=True, max_overlap=1).

## Accession numbers

Sequence data generated in this study will be submitted to the NCBI Sequence Read Archive upon manuscript acceptance. In addition, any file may be obtained from the corresponding author.

## Author contributions

BH, AF, and CS conducted wet lab experiments. SR, RW, and AW performed DL. BH, SR, and AW contributed to the data analysis. AR and AW designed the experiments and acquired funding. All authors contributed to the writing and editing of the manuscript and approved the final version prior to publication.

## Acknowledgments

The authors acknowledge and are appreciative of the intellectual contributions of Shyaron Poudel, Jeff Gaither, and the Arkansas High-Performance Computing Center.

## Supporting information

All code/additional information needed for the full reproduction of this study will be made available on GitHub upon acceptance of this manuscript. In addition, all files may be obtained from the corresponding author.

## Supporting information captions

**Fig S1** Capture-seq validation of RNA-seq data. (**a**) Heatmap of reference gene expression in RNA- and Capture-seq. (**b**) Heatmap of pathogen-induced gene expression.

**Fig S2** Selection of 201-bp bound (peak regions) and unbound sites (negative dataset) during model training.

**Fig S3** auROC and auPRC curves for soybean data-trained models. (**a**) auROC curve for GmWRKY30 CRNN. (**b**) auROC curve for GmRAV CRNN. (**c**) auPRC curve for GmWRKY30 CRNN. (**d**) auPRC curve for GmRAV CRNN.

**Fig S4** AmpDAP-seq data for GmMYB61 and GmWRKY2. (**a**) Heatmap of DAP peak binding within 1,000 bp of the TSS region. (**b**) Distribution of DAP peaks across genomic features.

**Fig S5** GRNs for defense-related TF families. (**a**) bHLH GRN. (**b**) ERF GRN. (**c**) MYB GRN. (**d**) NAC GRN. (**e**) RAV GRN. (**f**) WRKY GRN.

**Fig S6** Prioritization of target genes. (**a**) Density plots of indegree, cumulative outdegree, sum cumulative cosine, and mean |log_2_FC| (Mean |FC|) for target genes. Red polygons represent the upper quartile for each parameter. Furthermore, 254 targets were in the upper quarter for all four parameters. (**b**) Heatmap depicting the log_2_FC (FC) of prioritized target genes across both interactions.

**Table S1** RNA-seq mapping statistics.

**Table S2** Mapping statistics for DAP- and AmpDAP-seq.

**Data S1** DEGs from RNA-seq, corresponding co-expression cluster assignments, functional annotations, and target gene assignments.

**Data S2** Capture-seq gene list and expression data.

**Data S3** GmWRKY30 DAP-seq binding site data.

**Data S4** GmRAV DAP-seq binding site data.

**Data S5** GmMYB61 and GmWRKY2 AmpDAP-seq binding site data.

**Data S6** Arabidopsis generalization prediction accuracies.

